# Piezo2, a pressure sensitive channel is expressed in select neurons of the mouse brain: a putative mechanism for synchronizing neural networks by transducing intracranial pressure pulses

**DOI:** 10.1101/2020.03.24.006452

**Authors:** Jigong Wang, Owen P. Hamill

## Abstract

Piezo2 expression in the normal, young adult mouse brain was examined using an anti-PIEZO2 Ab generated against a C-terminal fragment of the human PIEZO2 protein. As a positive control for Ab staining of mouse neurons, the Ab was shown to stain the majority (~90%) of mouse dorsal root ganglia (DRG) neurons, consistent with recent in situ hybridization and transcriptomic studies that also indicate *Piezo2* gene expression in ~90% mouse DRG neurons. As a negative control and stringent test for specificity, the Ab failed to stain DRG satellite glial cells, which do not express *Piezo2* but rather its paralog, *Piezo1*. In slices of brains isolated from the same mice as the DRG, the Ab displayed high selectivity in staining only specific neuron types, including some pyramidal neurons in the neocortex and hippocampus, Purkinje cells in the cerebellar cortex, and most notably mitral cells within the olfactory bulb. Given the demonstrated role of *Piezo2* channels in peripheral neurons as a low-threshold pressure sensor (i.e., ≤ 5 mm Hg) critical for gentle touch, proprioception, and the regulation of breathing and blood pressure, its expression in select brain neurons has interesting implications. In particular, we propose that the pressure sensitive channel may provide specific brain neurons with an intrinsic resonance that acts to synchronize their firing with the normal pulsatile changes in intracranial pressure (ICP) associated with breathing and cardiac cycles. This novel mechanism could serve to increase the robustness of the respiration entrained oscillations that have been recorded in both rodent and human brains across widely distributed neuronal networks. The idea of a “global rhythm” within the brain has been mainly related to the effect of nasal airflow activating mechanosensitive neurons within the olfactory epithelium, which in turn synchronize, through direct synaptic connections, mitral neurons within the olfactory bulb and then through their projections, the activity of neural networks in other brain regions, including the hippocampus and neocortex. Our proposed, non-synaptic, intrinsic resonance mechanism for tracking pulsatile ICP changes would have the advantage that spatially separated brain networks could be globally synchronized effectively at the speed of sound.

## INTRODUCTION

In 2006, Kasuki Satoh and colleagues discovered that the rat ortholog of a human gene covering KIAA0233 (Nagase et al., 1998) was transcriptionally induced in cultured rat astrocytes treated with aggregates of beta-amyloid (Aβ) —a major component of senile plaques in Alzheimer’s disease (AD^+^) (Satoh et al., 2006). The gene was designated *Mib* (i.e., Membrane protein induced by Aβ) and predicted to encode a large membrane protein of >2000 amino acids that included >23 transmembrane domains. Most significantly, in situ hybridization (ISH) studies of human AD^+^ brains reported selective *hMib* expression in activated astrocytes surrounding Aβ-plaques, whereas in normal (i.e., AD−) brains, *hMib* expression was limited to neurons (Satoh et al., 2006). These key findings indicated that *Mib* expression is subject to an on-off switch that is cell-type sensitive and regulated by the surrounding brain conditions. In normal brains, the switch is turned on in some neurons, where presumably MIB performs a yet-to-be identified “homeostatic” function(s) important for neuronal function and/or survival. However, in AD^+^ brains, the switch is turned on in activated astrocytes, where MIB may take on a neuropathological role (Satoh et al., 2006). Since an over-expressed GFP-tagged Mib was localized to the endoplasmic reticulum (ER), it was suggested that Mib (also known as Fam38A) may function to regulate protein transport and signaling in the ER (Satoh et al., 2006) possibly involving integrin activation by Mib/Fam38A (McHugh et al., 2010).

In 2010, Bertrand Coste and colleagues significantly advanced the Mib story by using a siRNA knockdown screen to demonstrate that Mib forms a mechanically activated channel expressed in the plasma membrane of a mouse neuroblastoma cell line (Coste et al., 2010). Accordingly, Mib was re-designated Piezo1 to indicate a pressure-activated channel. Vertebrates express two Piezo family members, Piezo1 (Fam38A) and its paralog Piezo2 (Fam38B) that show differential expression in various mouse tissues including peripheral and central neural tissues (Coste et al., 2010). Most notably, in mouse DRG neurons *Piezo2* was very highly expressed compared with *Piezo1*, whereas in mouse brain *Piezo1* and *Piezo2* were expressed in equally low, but quantifiable levels (Coste et al., 2010) as one might expect if Piezo1 and Piezo2 were only expressed in very select brain neurons (Satoh et al., 2006). In the case of *Piezo1*, subsequent studies have confirmed *Piezo1* expression in the brain (Blumenthal et al., 2014; Pathak et al., et al., 2014; Zhang et al., 2014; Zeisel et al., 2015; Habib et al., 2016; Wang et al., 2019c). Most significantly, Blumenthal et al., (2014) studying rat hippocampal neuron-astrocyte interactions, proposed that the Piezo1 channel in neurons is able to sense and transduce the surface nanoroughness of astrocytes into trophic signals that maintain neuronal survival. However, in AD^+^ brains with the progressive build-up of mechanically stiff senile plaques, there is a detrimental increase in brain tissue nanoroughness that is transduced by Piezo1 into anti-survival signaling (Blumenthal et al., 2014; see also Velasco-Esterervez et al., 2018). Interestingly, other stressful conditions increase Piezo1 expression, including those occurring with normal aging, bacterial infection, brain ischemia or exposure to demyelinating agents. Furthermore, experimental over activation of Piezo1 channels using Yoda1, a Piezo1 specific agonist (Syeda et al., 2015) directly disrupts normal neuronal function and/or survival (Wang et al., 2019c; Velasco-Esterervez et al., 2019a, b).

In the case of Piezo2, most studies to date have focused on Piezo2 expression in the peripheral nervous system and its key mechanotransduction roles in gentle touch, proprioception, and the regulation of breathing and blood pressure (Coste et al., 2010; Ranade et al., 2014; Woo et al., 2014; 2015; Usoskin et al., 2015; Florez-Paz et al., 2016; Nonomura et al., 2017; Zeng et al., 2018). However, Venkata Sajja and colleagues have demonstrated, using RT-PCR and Western blotting using a commercial anti-Piezo2 antibody (Ab), that Piezo2 is expressed in rat neocortical and hippocampal tissue, and this expression is increased in response to repetitive mechanical (i.e., blast) injury (Sajja et al., 2014, Heyburn et al., 2019). These findings raise the possibility that a rapid mechanical activation of Piezo2 caused by concussion or mild traumatic brain injury (TBI) contributes to the rapid and often reversible disruptions in brain functions including consciousness and memory. Furthermore, the observed changes in Piezo2 expression caused by repetitive TBI may contribute to long term neurodegenerative processes and disorders that have been associated with TBI (Mortimer et al., 1991; Sajja et al., 2014, Nogueira, et al., 2016; Heyburn et al., 2019).

In this study, we use immunohistochemistry (IHC) and a custom generated anti-Piezo2 Ab against a C-termini fragment of the human PIEZO2, to investigate Piezo2 expression in the mouse brain, and in particular determine if Piezo2 is expressed in specific brain neurons. In brief, our results indicate that Piezo2 is expressed in some pyramidal neurons of the neocortex (NC) and in specific regions (CA3) of the hippocampus (HC), in Purkinje cells but not granule cells of the cerebellar cortex and perhaps most notably, in mitral cell of the olfactory bulb (OB). We propose that the activation of the Piezo2 cation channel by normal (< 10 mm Hg) or abnormal (> 25 mm Hg) intracranial pressures (ICP) may increase neural excitability thereby altering neural circuit activity and brain function. Furthermore, the occurrence of even very low level induced changes in Piezo2 activity may confer on key neurons an intrinsic membrane resonance that acts to synchronize their firing with the perennial pulsatile changes in ICP associated with breathing and cardiac cycles. In particular, this resonance could serve to increase the robustness of respiration entrained neural network oscillations observed within widespread brain regions —including the OB, HC and NC —and to date mostly thought to be driven by synaptic inputs from respiration-related peripheral afferents and/or efferent copy from brain stem respiratory nuclei (for reviews see Fontanini & Bauer, 2006; Heck et al., 2017; Tort et al., 2018).

## MATERIALS AND METHODS

### Isolation of mouse brains and dorsal root ganglia

All experimental protocols were approved by the Animal Care and Use Committee at the UTMB and are in accordance with the NIH Guide for the Care and Use of Laboratory Animals. Young adult (20–25 g, ~10 weeks old) male C57BL/6J or GAD67-GFP mice (Jackson Laboratory, Bar Harbor, Maine, USA) were deeply anesthetized with 5% isoflurane in oxygen, then after exposing the pleural cavity, perfused through the left ventricle first with cold heparinized saline aorta and then with 4% paraformaldehyde in 0.1 M phosphate-buffered saline (PBS). Their DRG and brains were rapidly removed and stored in the same fixative overnight. Subsequently, the brains and DRG were dehydrated through an ethanol series/xylene, embedded in paraffin and 10 μm sagittal slices cut. Human red blood cells were collected in the fixative, pelleted, dehydrated, and embedded in paraffin for slicing.

### Anti-Piezo2 antibody generation

The anti-PIEZO2 Ab used in this study was custom generated by Proteintech (Chicago, Ill) against a peptide fragment overexpressed from a cloned fragment of the human PIEZO2 (pF1KE0043) purchased from Kazusa DNA Research Institute (Chiba, Japan). This vector is suitable for expression of tag-free proteins in *E. coli* cells as well as in vitro protein-expression systems driven by T7 RNA polymerase. The antigen was a 188 amino acid peptide corresponding to amino acids 2440-2628 located at the C-terminus of the human PIEZO2 isoform (NP_071351.2), corresponding to amino acids 2512-2700 of the mouse PIEZO2 isoform (NP_001034574.4). The Ab was affinity purified using an antigen bound column. The human Piezo2 peptide displayed 84% amino acid identity (i.e., 158 out of 188 residue identity) with this mouse Piezo2 region. In comparison, the peptide showed only 34.6% amino acid identity (65 out of 188 residue identity) with the corresponding region (3226-3414) of the mouse Piezo1 isoform (NP_001032375.1). Significantly, and coincidently as it turned out, the antigen peptide we chose to generate our anti-PIEZO2 Ab, encompasses the shorter C-terminal peptide (75 amino acids and corresponding to amino acids 2446-2521 of the human PIEZO2 protein) that was used to generate the commercially available anti-PIEZO2 Ab (Atlas antibodies, HPA015986). This Atlas Ab was one of three antibodies used by the Human Protein Atlas group to study PIEZO2 expression in the human brain.^1^

Piezo2 immuno-reactivity was detected using the anti-Piezo2 Ab at a 1:600 dilution for 60 minutes. The brain and DRG tissues were processed using the Dako Autostainer. The secondary Ab used was biotinylated goat anti-rabbit IgG (Vector Labs, Burlington, CA, #BA-1000) at 1:200 for 30 minutes followed by Dako LSAB2 streptavidin-HRP(#SA-5704) for 30 minutes. Slides were developed with Dako DAB chromagen (#K3468) for 5 minutes and counterstained with Harris hematoxylin for one minute. Imaging was carried out under bright field Olympus Bx51 microscope with Olympus DP imaging software.

### Anti-Piezo2 antibody validation

In terms of Ab validation, the first question we asked was whether our anti-Piezo2 Ab could display the selectively to label Piezo2 in the brain rather than, or in addition to, its paralog, Piezo1? This was important since Piezo1 was first identified in rodent brain cells, and also shown to be expressed in specific neurons of the normal human brain (Satoh et al., 2006). Moreover, both Piezo1 and Piezo2 transcripts are expressed in equally low quantifiable levels in isolated mouse brain tissue (Coste et al., 2010). The second question was whether the Ab had the sensitivity to detect endogenous Piezo2 in a cellular pattern consistent with known endogenous Piezo2 gene expression in distinct murine cell types. This was also important in order to extend the published findings suggesting Piezo2 gene and protein expression in rat isolated neocortex and hippocampus tissue (Sajja et al., 2014; Heyburn et al., 2019).

To address both issues of selectivity and sensitivity, we took advantage of our recent study describing *Piezo1* and *Piezo2* expression in dorsal root ganglion (DRG) cells. In particular, the demonstration that Piezo2 transcripts are expressed in the large majority (~90%) of mouse DRG neurons of both small and large diameter but are undetectable in DRG satellite glial cells (SGC). (Wang et al., 2019a). *Piezo1* transcripts, although co-expressed with *Piezo2* in lower abundance and in a smaller proportion of small diameter DRG neurons, are strongly expressed in some SGCs (Wang et al., 2019a). The observations for *Piezo2* are consistent with recent ISH/RNAscope study that used the same *Piezo2* RNA probe (Zhang et al., 2019) and the case both *Piezo1* and *Piezo2* with two unbiased single cell RNA-sequencing studies (Usoskin et al., 2015; Sharma et al., 2020). Interestingly, sensory neurons in the mouse and human trigeminal ganglia (TG) also show similar high *Piezo2* expression (70 - 87%) (Szczot et al., 2017; Nickolls et al., 2020). However, in comparison, the very first studies identifying and characterizing *Piezo2* did indicate a lower proportion (20 −47%) of mouse DRG neurons expressing Piezo2 (Coste et al., 2010, Woo et al., 2014; Ranade et al., 2014).

The added advantage in choosing mouse DRG for Ab validation based on the known pattern of endogenous Piezo expression in mouse DRG neurons and SGCs (i.e., versus *Piezo2*^+^/Piezo1^+^ transfected Hek-293 cells) was that the DRG also provided an internal control against variations in Ab staining that might otherwise arise —due to differences in species, cell type and protocols used for tissue fixation and IHC — simply because the DRG and brains were isolated and fixed from the same mouse at the same time, and processed with identical IHC protocols. As it turned out, this internal control was important because whereas the anti-Piezo2 Ab stained the large majority of DRG neurons, it displayed extreme selectivity in staining very specific neuron types (e.g., Purkinje cells, mitral cells and CA3 neurons) and not staining more abundant other neuron types in the same brain regions (e.g., granule cells in the cerebellar cortex, olfactory bulb and hippocampus).

## RESULTS

### The anti-PIEZO2 antibody stains most mouse DRG neurons but no satellite glial cells

Figure 1A indicates that the anti-PIEZO2 Ab stained positive (i.e., brown) the majority of neurons in the DRG without staining the cell bodies of the satellite glial cells (SGCs) that ensheath the DRG neuron (i.e., blue stained cells). Only a minority (~10%) of DRG neurons were judged to show little or no staining (i.e., indicated by the asterisks) (Fig.1 B). For 10 DRG isolated from 2 mice, 1890 neurons were clearly stained by the Ab compared with only 110 that could be clearly identified as unstained. The Ab labeled DRG neurons ranged in diameter from 10 to 40 μm. This independence of soma size and thereby neuron type is consistent with most recent ISH and scRNA-seq studies (Usoskin te al., 2015; Zhang et al., 2019; Wang et al., 2019a; Sharma et al., 2020) that indicate both large and small diameter DRG neurons of most types show expression of *Piezo2* transcripts (but see Coste et al., 2010; Woo et al., 2014; Ranade et al., 2014) The lack of Ab staining of SGCs is also consistent with ISH results that indicate an absence of *Piezo2* transcripts (Ranade et al., 2014; Zhang et al., 2019; Wang et al., 2019a). The SGCs thereby provide a negative control for the anti-Piezo2 Ab as well as a stringent test for Ab specificity since SGCs express Piezo1 (Wang et al., 2019a). We also saw a similar lack of Ab staining of human RBCs that selectively express Piezo1 (Ma et al., 2018; data not shown).

**Figure 1.**
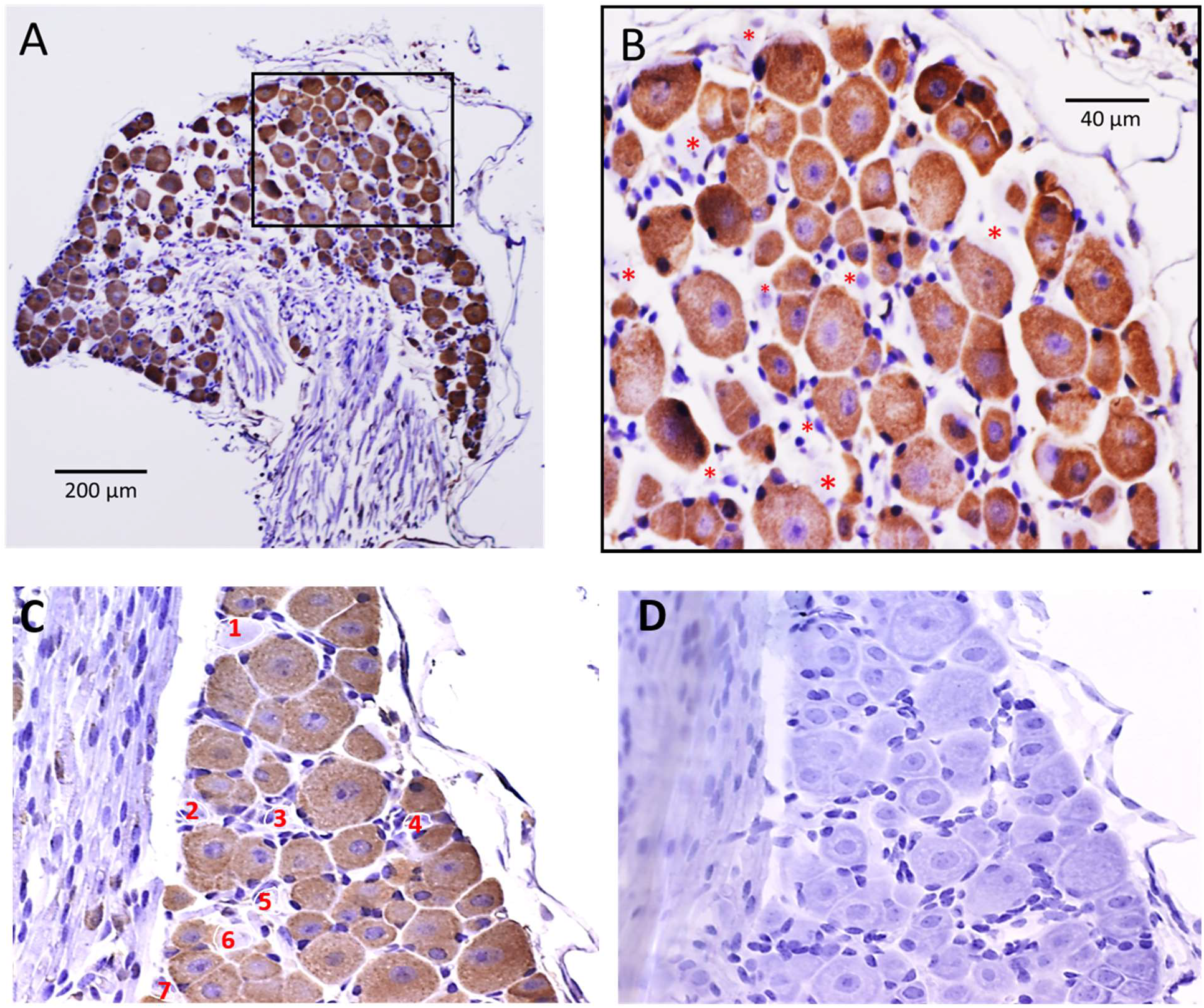
Immunohistochemical localization of Piezo2 in a mouse DRG slice. A: The anti-Piezo2 Ab stained the majority of DRG neurons independent of soma size (range 10-50 ◻m in diameter). B: Higher magnification image of the section enclosed within the square in 1A shows that of the ~ 90 neurons present, only 9 neurons (i.e., indicated by red asterisks) showed no Ab staining and these appeared to be smaller diameter neurons. The numerous small blue structures (i.e., stained by the nuclear stain hematoxylin) around the perimeter of each neuron represent the cell bodies of satellite glia cell (SGC) that includes their single cell nucleus and cytoplasmic processes that ensheath the neuron that showed no staining by the anti-Piezo2Ab. C: Another DRG slice from a different mouse showing a similar pattern of staining most DRG neurons but no SGCs. D: The same slice treated with the same IHC protocol except the anti-Piezo2 antibody was left out.

The DRG was further used to test for any possible nonspecific staining by the secondary Ab; a test achieved by leaving out the primary Ab, but otherwise completing the IHC protocol (c.f., Figs 1C &D). We did not perform a pre-adsorption test using the immunizing antigen since our Ab had already been affinity purified using an antigen-bound column. We recognize that neither test can rule out the possibility that our anti-Piezo2 Ab is not recognizing Piezo2 —alone or at all but – rather some off-target protein that possesses an immune-mimetic epitope. However, in this scenario the hypothetical off-target protein, but not the very closely related Piezo1, would need to share the immune-mimetic epitope. Moreover, the off-target protein would also need to share the same all-or-none pattern of Piezo2 gene expression in DRG neurons and SGCs. And as described below, display a highly selective expression in relatively few and very specific neuron types in the brain.

One test that might exclude an off-target protein, is to show the Ab labels in DRG extracts a single band with a MW consistent with Piezo2 (i.e., 318 kDa vs 286 kDa for Piezo1). However, this distinction may be problematic given that large proteins tend to run faster than predicted on gels depending upon the specific protein (Rath et al., 2013). Moreover, using WBs to validate Abs for IHC raises another caveat (also a common cited rationale) related to the specific tertiary structure of the immune epitope that the Ab recognizes. In our case, in an attempt to maximize Ab sensitivity we chose a C-termini peptide 2440-2628, based on the evidence that this forms part of the extracellular domain whose tertiary structure may contribute to Piezo2 pressure sensitivity (Taberner et al 2019; Wang et al., 2019b). Therefore, while this epitope \may be fully accessible for Ab binding in IHC given that the protein’s tertiary structure may be better preserved in the fixed and still membrane embedded state, it may be lost or concealed in the denatured globular form of the protein that is floating in solution, as in WB. The reverse situation may also apply where the epitope becomes more accessible in the denatured, globular and free solution form. An anti-Piezo2 Ab generated by Woo et al., 2014 did label a single band of > 270 kDa in WBs of Piezo2 transfected Hek cells and also labeled a lower percentage (<50%) of DRG neurons (Woo et al., 2014; Ranade et al., 2014). While the lower % was concordant with Piezo2 gene expression measured by the same group, it is discordant with more recent ISH and unbiased scRNA studies (see refs above).

To summarize, given the apparent concordance between Piezo2 protein and mRNA expression evidenced by our Ab and recent ISH and scRNA-seq studies, several validation criteria are met.

i. A demonstrated sensitivity and selectivity in staining neurons with confirmed endogenous levels of *Piezo2* gene expression in Piezo2+neurons and a lack of straining of *Piezo2*^−^ SGCs.
ii. The all-or-none nature of *Piezo2* gene expression in DRG neurons versus SCG provides a fairly stringent test against false negatives and false positives, respectively.
iii. The *Piezo1*+ SGC also provides a stringent test for lack of cross reactivity of the anti-Piezo2 Ab with its paralog Piezo1

### The anti-Piezo2 antibody stains pyramidal neurons in the mouse neocortex

The anti-Piezo2 Ab stained many cells throughout the grey matter of the neocortex, most notably cells spanning across the layer V-VI boundary (Figure 2A). Examination at high magnification indicated that the stained neurons were mostly pyramidal-like, as indicated by their triangular shaped cell body and unitary (apical) dendrite oriented perpendicular to the cortical layers and directed towards the pia (Figure 2B). In comparison, to the staining of pyramidal neurons, we saw no evidence of staining of multipolar cells that may have represented stellate interneurons or astrocytes.

**Figure 2.**
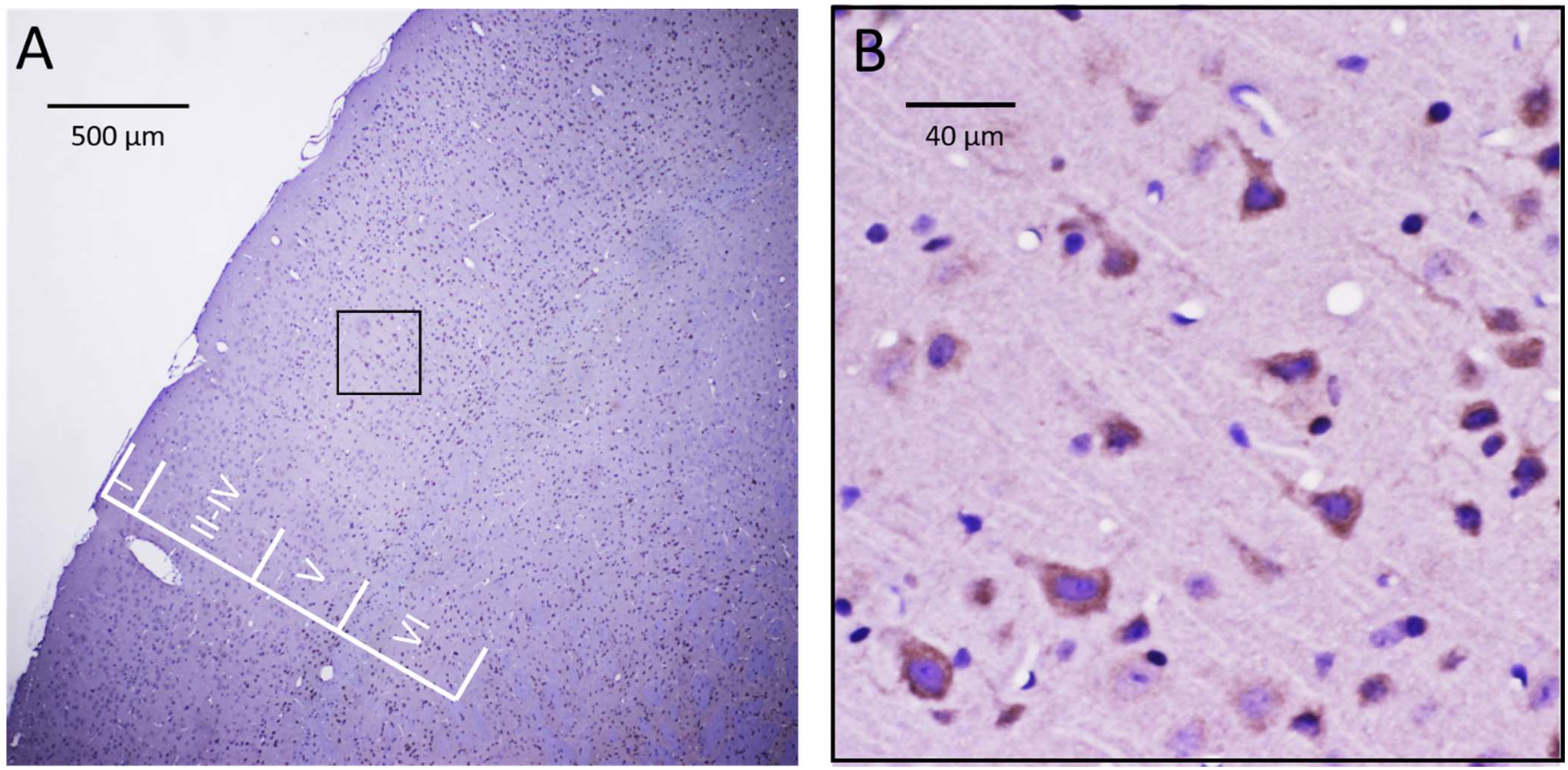
Immunohistochemical localization of Piezo2 in the mouse neocortex. A: The anti-Piezo2 Ab stained many cells throughout layers II-VI of the neocortex with minimal staining of cells in layer I. B: The selected region in A (i.e., within the black box) is shown at higher magnification and indicates that the most commonly stained cells displayed a pyramidal shaped cell body with a thick process directed towards the pia (i.e., consistent with the apical dendrite of a pyramidal neuron). In contrast, there was no clear evidence of the staining of multipolar cells indicative of stellate neurons or astrocytes.

### The anti-Piezo2 antibody stains neurons in specific regions of the hippocampus

The anti-Piezo2 Ab stained differentially neurons in specific regions of the hippocampus as designated in Figure 3A (dentate gyrus, CA1-4). Figure 3B shows a magnified section of the CA3 region in which at least 10 neurons out of ~ 35 neurons in the field showed staining by the anti-Piezo2 Ab. Figure 4 shows individual regions of the hippocampus at high magnification in which stained versus unstained cells were counted. In particular, figures 4A-C show representative regions of the dentate gyrus, in which 1,149 cells were recognized and counted. Of these cells only 16 cells (1.4%) were clearly stained by the Ab, and they were mostly spindle shaped cells located in the sub-granular layer, possibly reflecting interneurons. Similarly, in figure 4D showing the hilius/C4 regions, 72 neurons were counted but only 2 neurons stained positive (< 3%). These regions contrasted with the CA3 and CA2 regions (Figs, 4E-H), where 36% (50 cells out of 140) and 22% (15 cells out of 71) stained positive, respectively. On the other hand, in the CA1 region (Figs. I-L) only a relatively smaller percentage of neurons (~ 8%) were stained (i.e., 23 out of 305). Of the neurons in the CA regions that were Piezo2 positive most appeared to be pyramidal neurons as judged by their location and dendritic tree. A similar selective pattern of relatively higher proportional staining of neuron in the CA3-CA2 regions (75 of 240 cells, 30%) compared with dentate/CA4 (76 of 3,011, 2.5%) and CA1 (37 out of 540, 7%) regions was seen in hippocampi isolated from 2 different mouse brains.

**Figure 3.**
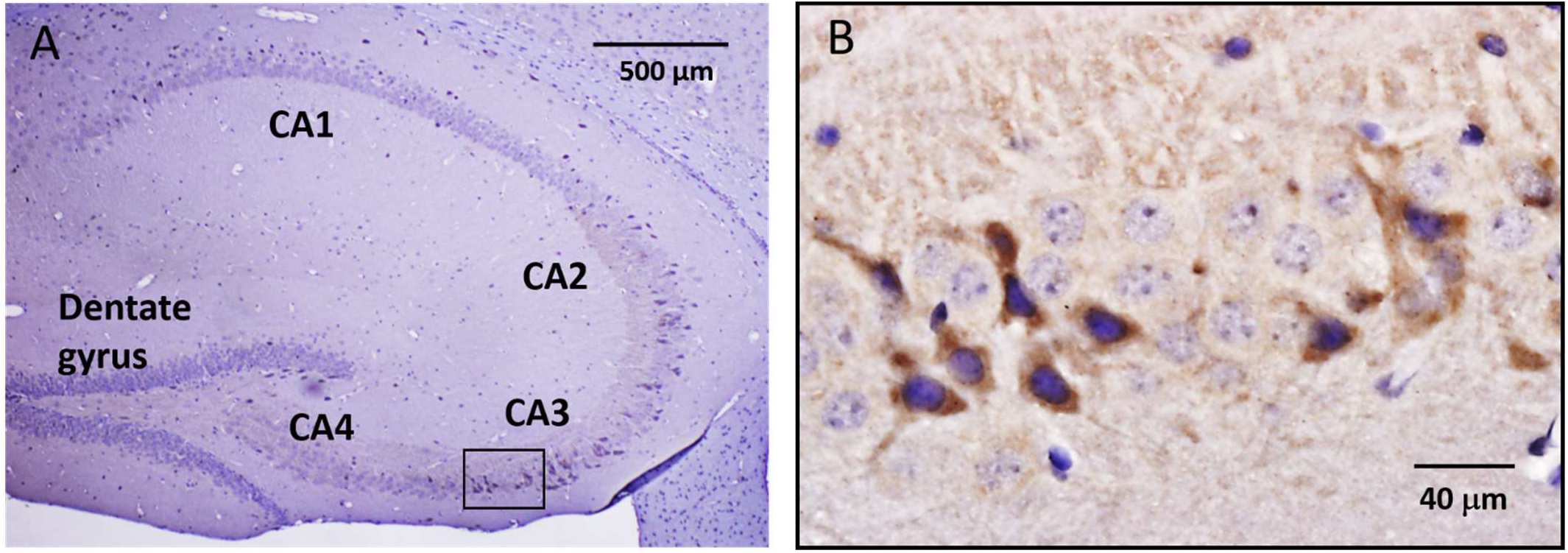
Immunohistochemical localization of Piezo2 in mouse hippocampus. A: Low magnification image of a hippocampal slice showing its distinct cellular regions that include the dentate gyrus (DG) and CA4-CA1 regions. B: The selected region in A (i.e., within the black box) that involves part of the CA3 region is shown at higher magnification and indicates at least 11 heavily stained cells that in some cases display a pyramidal shaped cell body with a single prominent process. Within the same region, at least 25 cells that were stained blue by hematoxylin showed no staining by the Ab. This all-or-none staining is at least consistent with idea of a stochastic on-off switch controlling Piezo2 expression.

**Figure 4.**
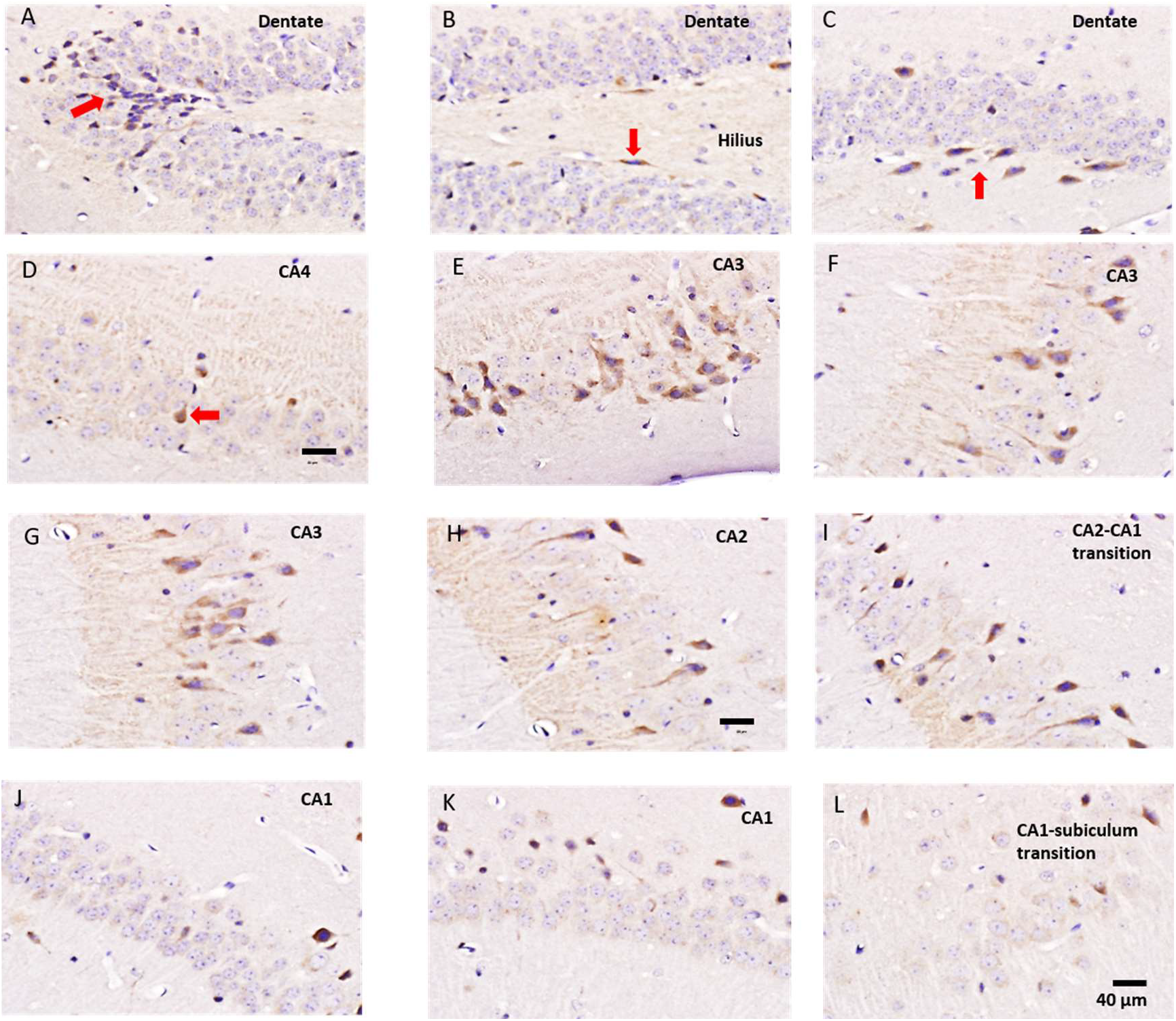
Differential anti-Piezo2 Ab staining of cells in different regions of the hippocampus examined at higher magnification. A: Shows the tip of the DG in which the majority of granule cells (i.e., round blue cells) are unstained but with a few cells at the very tip of the gyrus (red arrow) displaying some evidence of cytoplasmic staining. B: Shows the adjacent region of the DG in which 2-3 spindle shaped cells (one indicated by red arrow) located in the subgranular zone show some Piezo2 Ab staining. C: A continuing region further along the DG showing ~ 7 spindle shaped cells stained by the anti-Piezo2 Ab. D: Shows the C4 region with at least one cell darky stained cell (red arrow). E-G: Adjacent CA3 regions showing the highest density of Ab staining of neurons, many with unitary thick processes (i.e., apical dendrites) consistent with pyramidal neurons. H: CA2 region showing a lower density of stained neurons compared with the adjacent CA3 region. I: A region near the CA2-CA1 transition that displays a further reduced density of stained neuron. J-K: CA1 regions showing very few stained neurons. L: CA1 region transitioning into the subiculum that displayed no Ab-stained cells.

### The anti-Piezo2 antibody selectively stains mitral cells in the olfactory bulb

As indicated in figure 5A, that shows a low magnification image of the mouse olfactory bulb, the anti-Piezo2 Ab stained the neuropil of the OB whereas the hematoxylin stain delineated the circular glomeruli. In progressively higher magnified images (Figs. 5B-D), the strongest staining was of the mitral cells that form a distinct boundary cell layer separating the plexiform and granular cell layers, neither of which showed cell staining. The staining of the neuropil in the external plexiform layer presumably represents stained mitral cell dendrites that project to the glomeruli where they receive synaptic inputs from primary olfactory neurons. Tufted cells located in the external plexiform layer that also receive synapses from PON did not appear as clearly stained as the layer of mitral cells. As indicated in Fig. 5D, the most abundant cells of the OB, the granule cells were not stained by the Ab.

**Figure 5.**
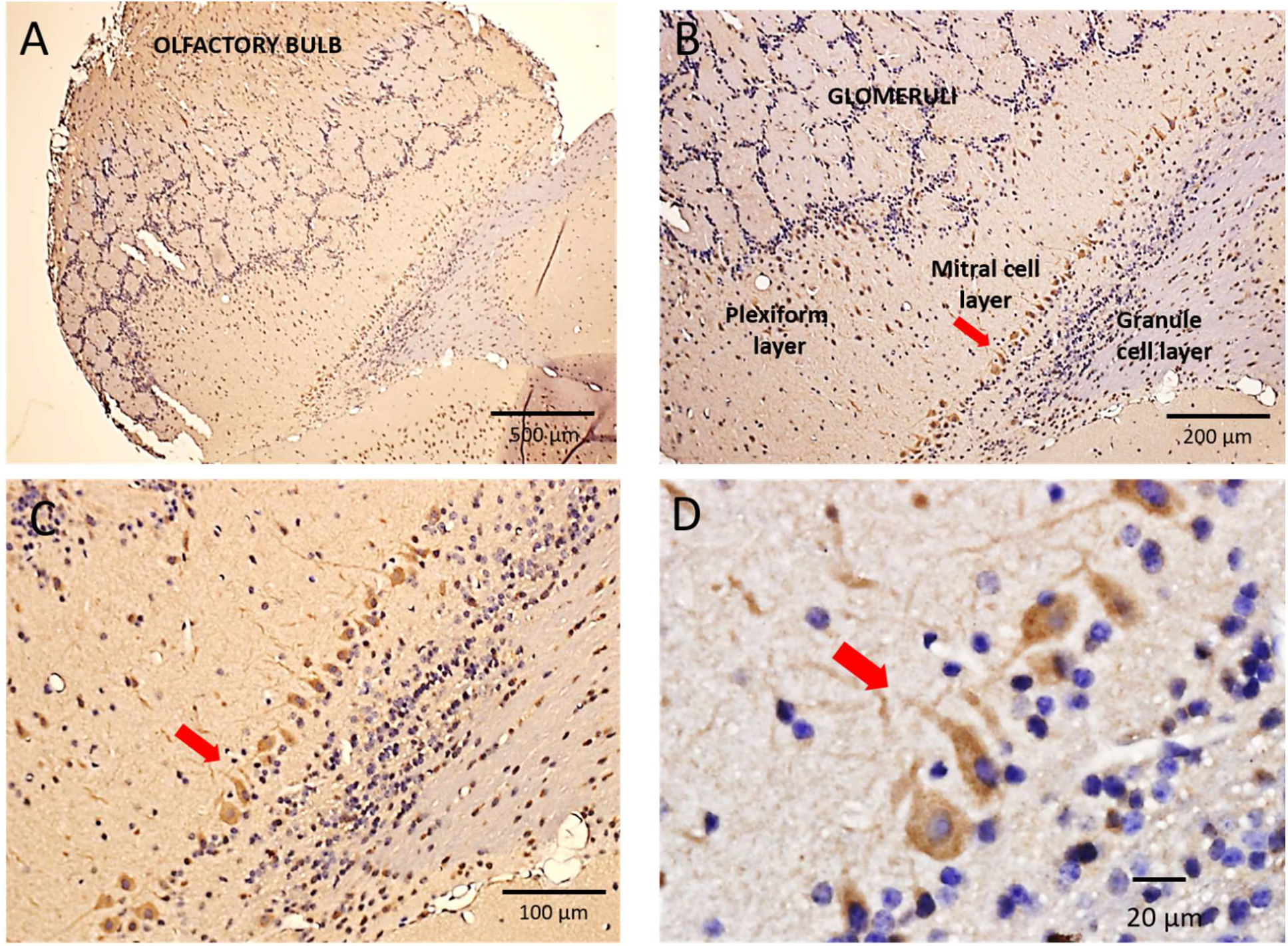
Immunohistochemical localization of Piezo2 in the mouse olfactory bulb. A: Shows a low magnification image of the mouse olfactory bulb (OB) with its characteristic circular/spherical glomeruli structures spanning across the OB. These glomeruli include the synaptic connections formed between primary olfactory nerve axons and mitral cell dendrites. B, C: Higher magnification images (10x and 20x objectives) of the same slice showing the uniformly stained layer of mitral cell bodies (red arrows) that separate the external plexiform layer from the internal plexiform and granule cell layers. The mitral cells represent the major projection neuron of the OB and project their axons to the piriform and entorhinal cortices as well as the amygdala. The external plexiform layer includes the primary and lateral dendrites of the mitral cells that extend into and throughout the plexiform layer to reach the glomeruli. Also within this layer are the cell bodies and dendrites of the tufted cells that did not appear stained. D: A still higher magnified image (60x objective) showing the morphology and staining of the mitral cells bodies and dendrites as well as the absence of staining of granule cells and the tufted cells in the granule cell and external plexiform layers, respectively.

### The anti-Piezo2 antibody stains Purkinje cells of the cerebellar cortex

The anti-Piezo2 Ab stained neurons located in highly specific regions of the cerebellar cortex (Fig. 6). Most notably there was consistent staining of the large cell body and branching dendrites of Purkinje cells that form a single cell layer separating the granular and molecular cell layers that showed no evidence of stained cell types. However, in addition to the staining of the Purkinje cells, some neurons embedded within the white matter of the cerebellum (arbor vitae) and presumably representing projection neurons within a deep cerebellar nucleus, were stained by the anti-Piezo2 Ab.

**Figure 6.**
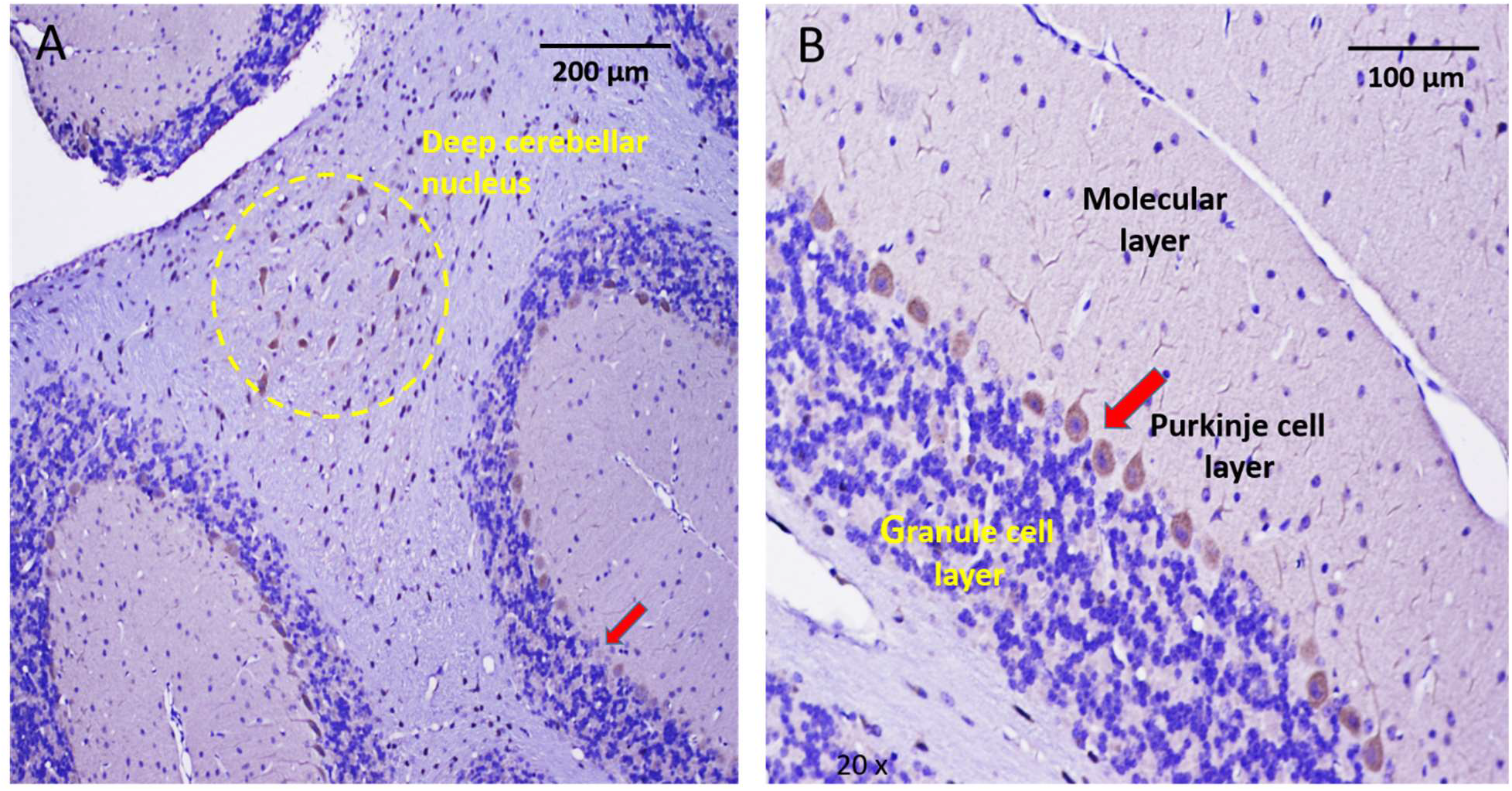
Immunohistochemical localization of Piezo2 in the cerebellum. A: Low magnification of a cerebellar slice indicating the stained Purkinje cell layer (red arrow) that separates the granule and molecular cell layers. The yellow dashed circle located within cerebellar white matter encloses a group of stained neurons presumably part of a deep cerebellar nucleus. B: Higher magnification of the cerebellar cortex showing the distinct line of Purkinje cells uniformly stained by the anti-Piezo2 Ab. In contrast to the Purkinje cells, the more numerous granular cells located within the granular cell layer were not stained by the Ab. Similarly, apart from Purkinje cell dendrites there was no pronounced Ab staining of cells in the molecular layer.

## DISCUSSION

This IHC study indicates that the low-threshold, pressure-activated channel Piezo2, is expressed in pyramidal neurons of the mouse neocortex and hippocampus, Purkinje cells of the cerebellar cortex and most notably, mitral cells of the olfactory bulb. This expression can be compared with PIEZO2 expression in human brain, as reported by the Human Protein Atlas (Uhlen et al., 2015).^2^ HPA is an Ab-based, proteomic-imaging analysis aimed at characterizing all proteins expressed in the human body. In the case of PIEZO2, one of the three HPA antibodies tested was generated against a C-terminus peptide (amino acids 2446-2521) that stained neurons in the human neocortex and hippocampus, and also cerebellar Purkinje cells (the human olfactory bulb was not examined). In the case of the other two antibodies one generated against an N-terminus peptide (amino acids 146-212) and the other against an internal region peptide (amino acids 1243-1275) — they did not stain neurons in any of the brain regions. However, the same antibodies also failed more often to stain cells in specific peripheral tissues, which may indicate a tissue type-dependent antibody sensitivity issue. In any case, the combination of our mouse Ab brain results with the HPA Ab result serves to increase the reliability of Piezo2 expression in the mammalian brain by i) adding an Ab that has been validated with known endogenous Piezo2 expression in DRG neurons ii) showing that two independently generated Abs display similar patterns of neuron staining in different species iii) providing added consistency with the literature that indicates Piezo2 expression in rodent brain measured by RT-PCR (Coste et al., 2010; Sajja et al., 2014) and Westerns (Heyburn et al., 2019). In addition, Zeisel et al., (2015)^3^ used quantitative single cell RNA sequencing (scRNA-seq) to identify mRNA transcripts in specific neurons in the mouse somatosensory (S1) cortex and the CA1-2 regions of the hippocampus. Their data reproduced in Figure 7, indicate *Piezo2* and *Piezo1* transcripts, in overall very low copy numbers, but in relatively higher numbers in specific S1 and CA1-2 pyramidal neuron types compared with interneurons. Interestingly, more recent scRNA-seq data released on the Allen Brain Atlas website^4^ indicates the highest *Piezo2* transcripts are detectable in select neurons of the mouse CA3 region, at least consistent with our IHC results.. Regarding the low copy numbers, especially when compared with the housekeeper gene Peptidyl-prolyl cis-trans isomerase B (Ppib), it may be that Piezo channels even when expressed at extremely low density can exert powerful influence on neuron excitability. This is exemplified by the demonstration that the activation of only a single Piezo-like channel is sufficient to trigger repetitive firing in neocortical or hippocampal pyramidal neurons (see Nikolaev et al., 2015).

**Figure 7.**
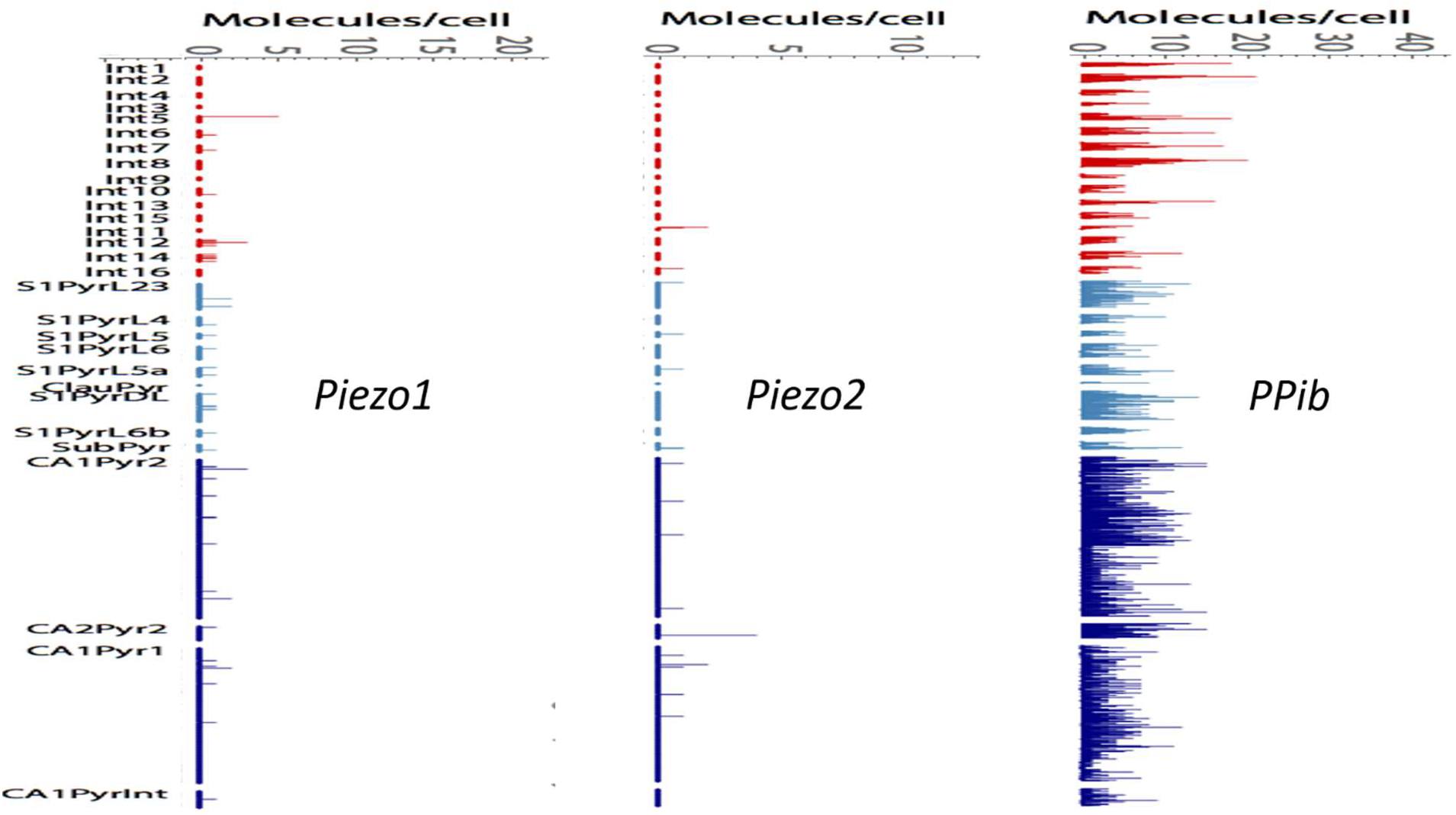
Quantitative single cell RNA-sequencing i(scRNA-seq) of single neurons isolated by papain dissociation from the somatosensory cortex (S1) and the hippocampal CA1-2 regions of mice (CD-1) of between 21 and 31 postnatal days and including both sexes. Data taken from Zeisel et al., (2015) (see http://linnarssonlab.org/cortex/). The vertical axis shows the genetically identified interneuron and S1 and CA1-2 pyramidal neuron types. The horizontal axis represent the single RNA transcript counts of Piezo1, Piezo2 and the housekeeper gene Ppib (Peptidyl-prolyl cis-trans isomerase B).

### Intracranial pressures in the brain that might activate the Piezo2 channel

Given the above evidence for pressure sensitive channels in brain neurons, the issue becomes the possible nature of the pressures that could activate the channels, and what might be the effects on brain function? Sajja and colleagues have begun to address the neuropathological case of traumatic brain injury (TBI) by demonstrating that blast forces alter Piezo2 channels expression, particularly in response to repetitive brain injuries (Sajja et al., 2014; Heyburn et al., 2019). Furthermore, TBI is just one of several neuropathological conditions including hydrocephalus, cerebral hemorrhage and brain tumors, that are known to elevate the normally low (< 10 mm Hg) baseline intracranial pressure (ICP) to higher levels (> 25 mm Hg) all with serious clinical consequences (Czosnyka & Pickard, 2004; Wagshul, et al., 2011; Andresen et al., 2015). Since Piezo2 can be activated by pressures as small as ≤ 5 mm Hg (Shin et al., 2018) the ICPs of 5 fold or more higher that develop under various neuropathological conditions, might be expected to induce abnormal Piezo2 channel activities and thereby contribute to deficits in brain function.

Perhaps the more interesting question is whether smaller changes in ICP (< 10 mm Hg) that occur in the healthy brain may also produce changes in Piezo2 channel activity and thereby alter brain function? Although the ICP in vertebrates is maintained at low mean baseline values of 0-10 mm Hg (Czosnyka & Pickard, 2004) it also undergoes repetitive, pulsatile changes, involving rapid (~200 ms) pulses with each heartbeat, as well as much slower (5-10 s) changes associated with shifts in venous blood and CSF between the brain and spinal cord during the breathing cycle (Czosnyka & Pickard, 2004; Dreha-Kulaczewsk et al., 2015; 2017). Moreover, specific volitional breathing practices —involving slow inspiration/expiration cycles and/or diaphragmatic vs thoracic breathing — performed to improve attention or reduce stress/anxiety, have been shown to cause even larger pulsatile changes in ICP (Aktas et al., 2019). This raises the intriguing possibility that the presence of Piezo2 in specific brain neurons allows them to transduce the amplified ICP changes, thereby altering neural network activity, EEG rhythms and causing the reported changes in brain functions (Moore et al., 2012; Zelano et al., 2016; Herrero et al., 2018).

### Respiration entrained oscillations occur in widespread neural networks of rodent and human brains

The idea that breathing can induce changes in neural network activity goes back nearly 80 years, with the discovery that nasal breathing in rodents causes rhythmic firing of mitral neurons within the olfactory bulb (OB) and neurons within the piriform cortex, independent of olfactory stimuli (Adrian, 1942). More recent studies have verified this discovery and shown that nasal breathing, not only entrains OB oscillations at the rodent’s breathing frequency (0.5-5 Hz), but also modulates the amplitude of higher frequency oscillations (80 - 120 Hz) in the OB as well as other downstream brain regions including the hippocampus (HC) and neocortex (NC). Moreover, these neural network oscillations have been linked to specific changes in rodent behaviors including whisking, memory formation/ consolidation as well as emotional (fear) responses (Fontanini et al., 2003; Yanowski et al., 2014; Nguyen Chi et al., 2014; Ito et al., 2014; Tsanov et al., 2014; Lockman et al., 2016; Zhong et al., 2017; Biskamp et al., 2017; Lockman & Tort, 2018; Moberly et al., 2018). Significantly, many of the observations related to respiration-locked oscillations seen in rodents, have been confirmed in humans using either intracranial EEG (iEEG) recording from epileptic patients or high density EEG recording from healthy humans (Zelano et al., 2016; Herrero et al., 2018; Piarulli et al., 2019; Perl et al., 2019).

Currently there are at least three, non-mutually exclusive, neuronal/biophysical mechanisms that may explain the widespread respiration synchronized brain oscillations (for reviews see Fontanini & Bower, 2006; Kay et al., 2009; Tort et al., 2018; Heck et al., 2019). The first, is a “peripheral afferent” mechanism dependent upon nasal air pressure changes activating mechanosensitive primary olfactory neurons (PON) within the nasal epithelium (Grosmaitre et al., 2007). These PON in turn activate, via their direct synaptic connections, mitral cells within the OB. The OB is then proposed to act as a “global clock” for other brain regions, synchronizing, via its synaptic connections, neural networks across widespread brain regions, including the HC and NC (Tort et al., 2018; Heck et al., 2019). Evidence supporting this mechanism, is that tracheotomy or ablation of the nasal epithelium (or removal of the OB) in rodents reduces the synchronized activity in the OB and/or in downstream brain regions (Adrian, 1942; Ravel & Pager, 1990; Fontanini et al., 2003; Phillips et al., 2012). Moreover, in both humans and rodents, air pressure pulses delivered to the nostrils can restore and/or alter the oscillation frequency (Lockman et al., 2016; Piarulli et al., 2018). The second mechanism depends upon “efferent copy” discharges from neurons in brain stem nuclei that regulate breathing (Kleinfeld et al., 2014; Yackle et al., 2017; Karalis & Sirota, 2018). Evidence for this mechanism, includes the finding that although tracheotomy or ablation of the nasal epithelium reduces the oscillations in rodents, they fail to completely block them (Fontanini et al., 2003; Karalis & Sirota, 2018). Moreover, in humans the oscillations are seen with mouth as well as with nasal breathing (Perl et al., 2019). Finally, it has been directly demonstrated that neurons in brain stem respiratory nuclei form connections with neurons in the locus coeruleus, which through their projections alter activity in higher brain regions (Yackle et al., 2017). The third mechanism, and the least investigated, proposes neurons possess an “intrinsic resonance” that by some unknown biophysical mechanism pre-tunes the neuron to fire at a frequency close to the respiratory frequency (Heck et al., 2019). The evidence supporting this mechanism, but also the 2nd mechanism, are that oscillations are retained even in the absence of nasal air flow or functional PONs, and during mouth instead of nasal breathing (Fontanini et al., 2003; Karalis & Sirota, 2018; Perl et al., 2019).

Here we propose a novel version of the third mechanism based on the presence of the Piezo2 channel in select brain neurons, most notably in mitral cells of the OB, and pyramidal neurons of the HC and NC. Specifically, we suggest that Piezo2 may confer on specific neurons an intrinsic resonance that acts to synchronize firing with ICP pulses. A schematic representing the three different mechanisms is shown in Figure 8 where axonal projections originating in the nasal epithelium (black arrow; mechanism I) and the medullary respiratory nuclei (green arrow; mechanism II) act —via their synaptic inputs (not represented) — to synchronize firing in the upstream brain regions including the OB, HC and prefrontal cortex. In mechanism III, the ICP pulse waveform is transduced by Piezo2 channel activity, which essentially results in almost simultaneous (i.e., at the speed or sound, 343 m/s) synchronization of all neural networks within wide spread brain regions. Although the focus here is on the breathing entrainment of memory circuit involving the OB, HC and PC (i.e., see Heck et al., 2019), the demonstrated presence of Piezo2 in cerebellar Purkinje cells in both human and mouse, may also play a role in the sniffing dependent activation of the human cerebellum (Sobel et al., 1998) and the highly correlated slow rate (~ 1 Hz) oscillations seen between mouse Purkinje cells (Cao et al., 2017).

**Figure 8.**
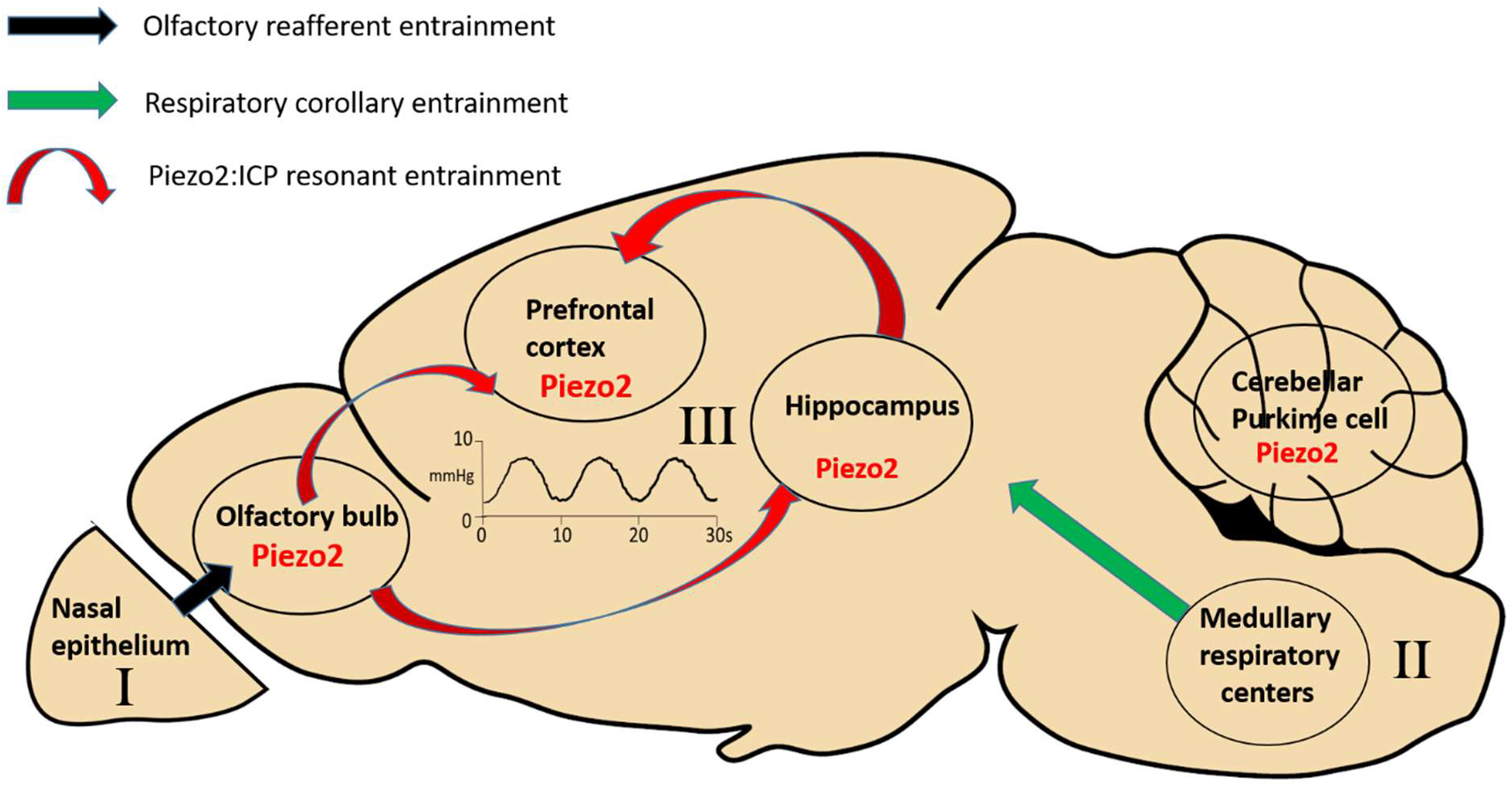
Schematic representation of three possible mechanisms by which respiration may cause entrainment of neuronal local field potentials in wide regions of the brain. In this case specifically, involving the olfactory bulb (OB), hippocampus and prefrontal cortex. The first mechanism involves respiratory olfactory reafferent (ROR) discharge and originates in the nasal epithelium where olfactory sensory neurons that also express mechanosensitive channels are able to detect the pressure changes associated with nasal airflow in the nostrils. Then through their direct synaptic connections (black arrow) with mitral cells and tufted cells in the OB cause respiratory entrainment of OB activity. In turn, the OB entrains neural activity in the hippocampus and prefrontal cortex through monosynaptic or polysynaptic connections (not shown) with these brain regions. The second mechanism, depends upon respiratory corollary discharge (RCD) from the neurons in the medullary respiratory nuclei that control the diaphragm muscles but also send efferent copy discharges to high brain regions (e.g., hippocampus and neocortex) by, as yet, unidentified synaptic pathways. The third mechanism, depends upon the pressure sensitive channel, Piezo2, shown here to be expressed in mitral cells of the OB and pyramidal cells of the hippocampus and neocortex. Piezo2, based on its demonstrated high pressure sensitivity, is proposed to transduce the intracranial pressure (ICP) waveform associated with the respiratory cycle, thereby providing a synchronizing clock for brain activity. This mechanism has the advantage over synaptic pathways in that the ICP pulse spreads throughout the brain faster (i.e., at 343 m/s the speed of sound) throughout the brain.

We envisage that the intrinsic resonance mechanism may work in concert to reinforce one or both of the synaptic driven mechanisms evident in rodents. However, it may be that the special features of human breathing have introduced selective pressures that add advantage to the intrinsic resonance mechanism. In particular, nasal breathing while obligatory in rodents is not in humans. Also, unlike in rodents where olfaction, sniffing and whisking are tightly linked and critical for their normal behavior and survival, olfaction is a relatively unimportant in humans. Moreover, humans are the only species capable of volitionally switching breathing patterns (e.g., to forced inspiration/expiration cycles) in order to promote changes in mood, cognition and emotions. Finally, humans are also unique in that they are able to consciously modulate through their breathing, both their heartbeat and emotions (Yasuma & Hayano, 2004; Mather & Thayer, 2018) possibly also via the associated changes ICP pulse frequency and waveform. Interestingly, one study of single unit recording from epileptic patients indicated that a higher proportion of cells in the hippocampus and amygdala were modulated by the cardiac cycle compared with the respiratory cycle (Frysinger & Harper, 1989).

From a mechanistic point of view, transducing the perennial and life spanning ICP pulses into membrane electrical oscillations, could act as a more robust/reliable clock mechanism, given its faster influence (i.e., at 343 m/s) compared with slower synaptic and axonal transmission. It would also be energy efficient by being driven by the indispensable cardio-respiratory cycles, as well as being less vulnerable to disruption compared with timing dependent on complex polysynaptic pathways. Finally, in a similar way that the various EEG recorded brain rhythms were considered an epiphenomena until their demonstrated key roles in information processing within the brain (Buzsaki, 2006) it will be interesting if ICP rhythms play an even more primary role by synchronizing the neural network activity that produce the EEG recorded electrical rhythms.

### Direct evidence for functional mechanotransduction in the brain

Despite the very early demonstration by Angelo Mosso (est. 1875) that changes in brain pressure, as measured by plethysmography, could be correlated with changes in human brain function (reviewed by Zago et al., 2009). Subsequent observations of mechanically induced changes in brain neuron activity — as when recording electrodes presses on brain tissue have — been more often considered as arising from non-specific or artefactual effects and not studied apart from their downstream consequences (see Ross et al., 2016). Similarly, in EEG recordings, direct mechanical disturbances associated with cardiac or breathing cycles have more often been seen as artefacts that need to be removed (e.g., Urigüen & Garcia-Zapirain 2015). Indeed, the observation that the respiration locked oscillations in human EEG recordings can be localized to cortical gray matter, rather than white matter or CSF containing compartments, has been used as evidence against mechanical artefacts (Herrero et al., 2018). However, the expression of Piezo2 in neuronal cell bodies that make up the gray matter could also explain this localization.

Interestingly, it is only in the last 10 years that the first direct and compelling evidence of significant mechanotransduction within the brain has come from a quite different research approach by William Tyler and colleagues who have demonstrated that transcranial focused ultrasound (tFUS) reversibly alters cortical circuit activity, EEG rhythms and behavior in both rodents and humans (Tyler, 2012; Tyler et a al., 2018). Moreover, Piezo1 channels have been directly implicated in mediating these tFUS effects (Prieto et al., 2018; Qiu et al., 2019). Somewhat surprising, there have been minimal attempts to directly study these channels in neurons in brain slices using patch clamp recording, the notable exception being a cell-attached patch clamp study that recorded single mechanically-gated cation channel currents in both neocortical and hippocampal pyramidal cells (Nikolaev et al., 2015). Significantly, the channel currents occurred spontaneously at a low frequency in the absence of applied suction and were increased in frequency with applied suction. Moreover, when activated by suction pulses of 20-30 mm Hg, the single channel currents occurred as bursts of rapid opening and closing that were sufficient to trigger repetitive firing in the whole cell (Nikolaev et al., 2015). Clearly a future priority will be to identify the molecules that form these channels, as well as characterize with patch recording the channel activity in the Piezo2 expressing neurons identified here.

## Conclusions

Finally, it is interesting to compare the differences in function between the peripheral baroreceptor reflex mediated by Piezo channels (Zeng et al., 2018) and the possible central baroreception function as proposed here. In the case of peripheral baroreception, the primary function is to ensure adequate blood supply to the body and brain, by regulating arterial blood pressure (Stocker et al., 2019 and refs therein). Peripheral baroreceptors achieve this by increasing or decreasing the sensory neuron discharge around a mean arterial pressure (MAP) set point of ~80 mm Hg, within a normal MAP range of around 40 to 180 mmHg. At the same time, responding dynamically to the pulsatile AP changes associated with each heartbeat. Therefore, their key function is to fine tune the afferent impulse activity to the brain as the sensory arm of the central mediated baroreceptor reflex. To perform this function, the baroreceptors need to display not only a high dynamic sensitivity to pulsatile pressure changes but also an ability to finely tune impulse discharge for small pressure changes (< 10 mmHg) over a broad dynamic pressure range. This second function may only be achieved by a relatively high expression/density of functional channels that are available to undergo calibrated activation over the MAP pressure range of 40 to 180 mm Hg. In contrast, the brain baroreceptor function we propose here may function quite differently, since ICP is typically maintained at very low values (e.g., <10 mm Hg). Moreover, cerebral perfusion pressure (CPP) and not ICP, is the most critical pressure for normal healthy brain function, which is mainly determined by the MAP (i.e., CPP = MAP – ICP). Instead of a homeostatic role, we hypothesize that ICP and Piezo2, acting together serve as a global clock mechanism for the brain, perhaps analogous to a system clock in a computer (see Tort et al., 2018). To achieve this function, ICP activation of Piezo2 activation should <u>not</u> normally evoke neural discharge (the exception being with pathological and elevated ICPs) as is required for peripheral baroreceptors, but rather modulate the membrane potential in a rhythmic fashion and thereby entrain or modulate the amplitude of neural circuit oscillations. In this case, a critical requirement for Piezo channels as brain baroreceptors may be a relatively low expression in select neurons.

## AUTHOR CONTRIBUTIONS

OH conceived the study, analyzed and interpreted the IHC experiments. JW surgically isolated the mouse brains and DRG. OH wrote the manuscript. All authors read and approved the final manuscript.

## ACKNOWLEDGMENTS

We thank Dr. Jin Mo Chung for providing financial support and Drs. Daniel Jupiter; Andrzej Kudlicki and Thomas Green for helpful discussions. We also appreciate the assistance from the UTMB Pathology core facility, and in particular Kerry Graves for preparing the brains and DRG slices for IHC. We also thank Kenneth Escobar for providing the microscopy facilities.

## Footnotes

Footnote 1: https://www.proteinatlas.org/ENSG00000154864-Piezo2/tissue/cerebellum

Footnote 2. https://www.proteinatlas.org/search/Piezo2

Footnote 3. http://linnarssonlab.org/cortex/

Footnote 4. http://celltypes.brain-map.org/rnaseq/mouse/cortex-and-hippocampus/

## REFERENCES

Adrian, E. D. (1942). Olfactory reactions in the brain of the hedgehog. J. Physiol. 100, 459–473.

Aktas, G., Kollmeier, J.M., Joseph, A.A., Merboldt, K.D., Ludwig, H.C., Gärtner, J., et al. (2019). Spinal CSF flow in response to forced thoracic and abdominal respiration. Fluids Barriers CNS. 16, 10. doi: 10.1186/s12987-019-0130-0.

Andresen, M., Hadi, A., Petersen, L.G., and Juhler, M. (2015). Effect of postural changes on ICP in healthy and ill subjects. Acta Neurochir (Wien). 157(1):109–13. doi: 10.1007/s00701-014-2250-2. Epub 2014 Oct 14.

Biskamp, J., Bartos, M., and Sauer, J.F. (2017). Organization of prefrontal network activity by respiration-related oscillations. Sci Rep. 7,45508. doi: 10.1038/srep45508.

Blumenthal, N.R., Hermanson, O., Heimrich, B., and Prasad Shastri, V. (2014). Stochastic nanoroughness modulates neuron–astrocyte interactions and function via mechanosensing cation channels. Proc. Nat. Acad. Sci. USA. 111, 16124–16129.

Bouchard, S., Bernier, F., Boivin, E., Morin, B., and Robillard, G.(2012). Using biofeedback while immersed in a stressful videogame increases the effectiveness of stress management skills in soldiers. PLoSOne. 7, e36169. doi: 10.1371/journal.pone.0036169. Epub 2012 Apr 27.

Buzsaki, G. (2006). Rhythms of the brain. Oxford: Oxford University Press.

Cao, Y., Liu, Y., Jaeger, D., and Heck, D.H. (2017). Cerebellar Purkinje Cells Generate Highly Correlated Spontaneous Slow-Rate Fluctuations. Front Neural Circuits. 11:67. doi: 10.3389/fncir.2017.00067. eCollection

Coste, B., Mathur, J., Schmidt, M., Taryn, J., Earley, T.J., Ranade, S., et al. (2010). Piezo1 and Piezo2 are essential components of distinct mechanically activated cation channels. Science 330, 55–60.

Czosnyka, M., and Pickard, J. (2004). Monitoring and interpretation of intracranial pressure. J Neurol Neurosurg Psychiatry. 75, 813–821.doi: 10.1136/jnnp.2003.033126

Dreha-Kulaczewski, S., Joseph, A.A., Merboldt, K.D., Ludwig, H.C., Gärtner, J., and Frahm, J. (2015). Inspiration is the major regulator of human CSF flow. J Neurosci. 35, 2485–2491. doi: 10.1523/JNEUROSCI.3246-14.2015. PMID: 25673843.

Dreha-Kulaczewski, S., Joseph, A.A., Merboldt, K.D., Ludwig, H.C., Gärtner, J., and Frahm, J. (2017). Identification of the Upward Movement of Human CSF In Vivo and its Relation to the Brain Venous System. J Neurosci. 37, 2395–2402. doi: 10.1523/JNEUROSCI.2754-16.2017. Epub 2017 Jan 30. PMID: 28137972.

Florez-Paz, D., Bali, K.K., Kuner, R., and Gomis, A. (2016). A critical role for Piezo2 channels in the mechanotransduction of mouse proprioceptive neurons. Sci. Rep. 6, 25923.

Fontanini, A., and Bower, J.M. (2006). Slow-waves in the olfactory system: an olfactory perspective on cortical rhythms. Trends Neurosci. 29, 429–437. Epub 2006 Jul 13.

Fontanini, A., Spano, P.F., and Bower, J.M. (2003). Ketamine-Xylazine-Induced Slow (< 1.5 Hz) Oscillations in the Rat Piriform (Olfactory) Cortex Are Functionally Correlated with Respiration. J. Neurosci. 23, 7993–8001; DOI: https://doi.org/10.1523/JNEUROSCI.23-22-07993.2003

Frysinger, R.C.,and Harper, R.M. (1989). Cardiac and respiratory correlations with unit discharge in human amygdala and hippocampus. Electroencephalogr Clin Neurophysiol. 72, 463–470.

Greitz, D., Wirestam, R., Franc, A., Nordell, B., Thomsen, C., and Ståhlberg, F. (1992). Pulsatile brain movement and associated hydrodynamics studied by magnetic resonance phase imaging. The Monro-Kellie doctrine revisited. Neuroradiol. 34, 370–80.

Grosmaitre, X., Santarelli, L.C., Tan, J., Luo, M., and Ma, M. (2007). Dual functions of mammalian olfactory sensory neurons as odor detectors and mechanical sensors. Nat Neurosci. 10, 348–54. Epub 2007 Feb 18.

GTEx Consortium. (2015). The Genotype-Tissue Expression (GTEx) pilot analysis: multitissue gene regulation in humans. Science. 348, 648–660.

Habib, N., Li, Y., Heidenreich, M., Swiech, L., Avraham-Davidi, I., Trombetta, J.J., et al. (2016). Div-Seq: Single-nucleus RNA-Seq reveals dynamics of rare adult newborn neurons. Science. 353, 925–8. doi: 10.1126/science.aad7038.

Heck, D.H., Kozma, R., Kay, L.M. (2019). The rhythm of memory: how breathing shapes memory function. J Neurophysiol. 122, 563–571. doi: 10.1152/jn.00200.2019.

Heck, D.H., McAfee, S.S., Liu, Y., Babajani-Feremi, A., Rezaie, R., Freeman, W.J. et al. (2017). Breathing as a Fundamental Rhythm of Brain Function. Front Neural Circuits. 12;10:115. doi: 10.3389/fncir.2016.00115. eCollection 2016.

Herrero, J.L., Khuvis, S., Yeagle, E., Cerf, M., and Mehta, A.D. (2018). Breathing above the brain stem: volitional control and attentional modulation in humans. J. Neurophysiol. 119, 145–159. Published online 2017 Sep 27. doi: 10.1152/jn.00551.2017

Heyburn, L., Abutarboush, R., Goodrich, S., Urioste, R., Batuure, A., Statz, J. et al. (2019). Repeated Low-Level Blast Overpressure Leads to Endovascular Disruption and Alterations in TDP-43 and Piezo2 in a Rat Model of Blast TBI. Front Neurol. 766. doi: 10.3389/fneur.2019.00766.

Ito, J., Roy, S., Liu, Y., Cao, Y., Fletcher, M., Lu, L., et al. (2014). Whisker barrel cortex delta oscillations and gamma power in the awake mouse are linked to respiration. Nat. Commun. 5:3572, DOI: 10.1038/ncomms4572

Karalis, N and Sirota, A. (2018). Breathing coordinates limbic network dynamics underlying memory consolidation. BioRXiv. doi: https://doi.org/10.1101/392530.

Kay, L.M., Beshel, J., Brea, J., Martin, C., Rojas-Líbano, D., and Kopell, N. (2009). Olfactory oscillations: the what, how and what for. Trends Neurosci. 32, 207–214. doi: 10.1016/j.tins.2008.11.008. Epub 2009 Feb 23.

Kleinfeld, D., Deschênes, M., Wang, F., and Moore, J.D. (2014) More than a rhythm of life: breathing as a binder of orofacial sensation. Nat Neurosci. May;17(5):647–51. doi: 10.1038/nn.3693. Epub 2014 Apr 25.

Lockmann, A.L., Laplagne, D.A., Leão RN, and Tort, A.B. (2016). A Respiration-Coupled Rhythm in the Rat Hippocampus Independent of Theta and Slow Oscillations. J Neurosci. 36, 5338–52. doi: 10.1523/JNEUROSCI.3452-15.2016.

Lockmann, A.L.V., and Tort, A.B.L. (2018). Nasal respiration entrains delta-frequency oscillations in the prefrontal cortex and hippocampus of rodents. Brain Struct Funct. 223, 1–3. doi: 10.1007/s00429-017-1573-1. Epub 2017 Dec 8.

Ma, S., Cahalan, S., LaMonte, G., Grubaugh, N. D., Zeng, W., Murth, S. E., et al. (2018). Common PIEZO1 allele in african populations causes RBC dehydration and attenuates plasmodium infection. Cell. 173, 443–455. doi: 10.1016/j.cell.2018.02.047

Mather. M., and Thayer, J. (2018). How heart rate variability affects emotion regulation brain networks. Curr Opin Behav Sci. 19, 98–104. doi: 10.1016/j.cobeha.2017.12.017.

McHugh, B.J., Buttery, R., Lad, Y., Banks, S., and Haslett, C. (2010). Integrin activation by Fam38A uses a novel mechanism of R-Ras targeting to the endoplasmic reticulum. J. Cell Sci. 123, 51–61.

Moberly A.H., Schreck, M., Bhattarai, J.P., Zweifel, L.S., Luo, W., and Ma, M. (2018). Olfactory inputs modulate respiration-related rhythmic activity in the prefrontal cortex and freezing behavior. Nat Commun. 9, 1528. doi: 10.1038/s41467-018-03988-1.

Moore, A., Gruber, T., Derose, J., and Malinowski P. (2012). Regular, brief mindfulness meditation practice improves electrophysiological markers of attentional control. Front Hum Neurosci. 6,18. doi: 10.3389/fnhum.2012.00018. eCollection 2012.

Mortimer, J.A., van Duijn, C.M., Chandra, V., et al. (1991). Head trauma as a risk factor for Alzheimer’s disease: a collaborative re-analysis of case-control studies. EURODEM Risk Factors Research Group. Int. J. Epidemiol. 20, S28–35.

Nagase, T., Seki, N., Ishikawa, K., Ohira, M., Kawarabayasi, Y., Ohara, O., et al. (1996). Prediction of the coding sequences of unidentified human genes: VI. The coding sequences of 80 new genes (KIAA0201-KIAA0280) deduced by analysis of cDNA clones from cell line KG-1 and brain. DNA Res. 3, 321–329.

Nguyen Chi, V., Müller, C., Wolfenstetter, T., Yanovsky, Y., Draguhn, A., Tort, A.B., et al. (2016). Hippocampal Respiration-Driven Rhythm Distinct from Theta Oscillations in Awake Mice. J. Neurosci. 36, 162–177. doi: 10.1523/JNEUROSCI.2848-15.2016

Nickolls, A.R., Lee, M.M., Espinoza, D.F., Szczot, M., Lam, R.M., Wang, Q., et al., (2020). Transcriptional Programming of Human Mechanosensory Neuron Subtypes from Pluripotent Stem Cells. Cell Rep. 30(3):932–946..

Nikolaev, Y.A, Dosen, P.J. Laver, D.R. Van Helden, D.F. and Hamill, O.P. (2015). Single mechanically-gated cation channel currents can trigger action potentials in neocortical and hippocampal pyramidal neurons. Brain Res. 1608, 1–13.

Nogueira, M. L., Epelbaum, S., Steyaert, J.M., Dubois, B., Schwartz, L. (2016). Mechanical stress models of AD pathology. Alzheimer’s & Dementia. 12, 324–333.

Nonomura, K., Woo, S.H., Chang, R.B., Gillich, A., Qiu, Z., Francisco, A.G., et al. (2017). Piezo2 senses airway stretch and mediates lung inflation-induced apnoea. Nature. 541,176–181. doi: 10.1038/nature20793. Epub 2016 Dec 21.

Pathak, M.M., Nourse, J.L., Tran, T., Hwe, J., Arulmoli, J., Le, D.T., et al (2014). Stretch-activated ion channel Piezo1 directs lineage choice in human neural stem cells. Proc Natl Acad Sci U S A. 111, 16148–16153. doi: 10.1073/pnas.1409802111.

Perl, O., Ravia, A. Rubinson, M., Eisen, A., Soroka, T, Mor, N., et al., (2019). Human non-olfactory cognition phase-locked with inhalation. Nat Hum Behav. 3, 501–512. doi: 10.1038/s41562-019-0556-z. Epub 2019 Mar 11.

Phillips, M.E., Sachdev, R.N., Willhite, D.C., and Shepherd, G.M. (2012). Respiration drives network activity and modulates synaptic and circuit processing of lateral inhibition in the olfactory bulb. J. Neurosci. 32, 85–98. doi: 10.1523/JNEUROSCI.4278-11.2012.

Piarulli, A., Zaccaro, A., Laurino, M., Menicucci, D., De Vito, A., Bruschini, L., et al. (2018). Ultra-slow mechanical stimulation of olfactory epithelium modulates consciousness by slowing cerebral rhythms in humans. Sci Rep. 26, 6581. doi: 10.1038/s41598-018-24924-9.

Prieto ML; Firouzi K; Khuri-Yakub BT; Maduke M. (2018). Activation of Piezo1 but Not NaV1.2 Channels by Ultrasound at 43 MHz. Ultrasound in Medicine & Biology. 44(6):1217–1232, 2018 06.

Qiu, Z., Guo, J., Kala, S., Zhu, J., Xian, Q., Qiu, W., et al., (2019). The Mechanosensitive Ion Channel Piezo1 Significantly Mediates In Vitro Ultrasonic Stimulation of Neurons. iScience. 21:448–457.

Ranade, S.S., Woo, S.H., Dubin, A.E., Moshourab, R.A., Wetzel, C., Petrus, M., et al. (2014). Piezo2 is the major transducer of mechanical forces for touch sensation in mice. Nature, 516, 121–125.

Rath, A., Cunningham, F., Deber, C.M. (2013). Acrylamide concentration determines the direction and magnitude of helical membrane protein gel shifts. Proc. Natl. Acad. Sci. U S A. 110, 15668–15673.

Ravel N, and Pager J (1990). Respiratory patterning of the rat olfactory bulb unit activity: nasal versus tracheal breathing. Neurosci Lett.115(2-3):213–218.

Ross, A.E., Nguyen. M.D., Privman, E., and Venton, B.J. (2014). Mechanical stimulation evokes rapid increases in extracellular adenosine concentration in the prefrontal cortex. J. Neurochem. 130, 50–60. doi: 10.1111/jnc.12711. Epub 2014 Apr 2.

Sajja, V.S.S.S., Ereifej, E.S., and VandeVord, P.J. (2014). Hippocampal vulnerability and subacute response following varied blast magnitudes. Neurosci Lett. 570, 33–37.

Satoh, K., Hata, M, Takahara, S, Tsuzaki, H, Yokota, H, Akatsu, H, et al. (2006). A novel membrane protein, encoded by the gene covering KIAA0233, is transcriptionally induced in senile plaque-associated astrocytes. Brain Res. 1108, 19–27.

Sharma, N., Flaherty, K., Lezgiyeva, K., Wagner, D.E., Klein, A.M., and Ginty, D.D. (2020). The emergence of transcriptional identity in somatosensory neurons. Nature. 577, 392–398.

Shin, K.C., Park, H.J., Kim, J.G., Lee, I.H., Cho, H., Park, C., et al. (2019). The Piezo2 ion channel is mechanically activated by low-threshold positive pressure. Sci Rep. 9, 6446.

Sobel, N., Prabhakaran,V., Hartley, C.A., Desmond, J.E., Zhao, Z., Glover, G.H., et al. (1998). Odorant-Induced and Sniff-Induced Activation in the Cerebellum of the Human. J. Neurosc. 18, 8990–9001. 10.1523/JNEUROSCI.18-21-08990.1998.

Stocker, S.D, Sved, A.F., and Andresen, M.C. (2019) Missing pieces of the Piezo1/Piezo2 baroreceptor hypothesis: an autonomic perspective. J Neurophysiol. 122, 1207–1212. doi: 10.1152/jn.00315.2019. Epub 2019 Jul 17.

Syeda, R., Xu, J., Dubin, A. E., Coste, B., Mathur, J., Huynh, T., et al. (2015). Chemical activation of the mechanotransduction channel Piezo1. eLife 4:e07369.

Szczot, M., Pogorzala, L.A., Solinski, H.J., Young, L., Yee, P., Le Pichon, C.E., et al. (2017). Cell-Type-Specific Splicing of Piezo2 Regulates Mechanotransduction. Cell Rep. 21, 2760–2771.

Taberner, FJ., Prato, V., Schaefer, I., Schrenk-Siemens, K., Heppenstall, P.A. and Lechner, S.G. (2019). Structure-guided examination of the mechanogating mechanism of PIEZO2. Proc Natl Acad Sci U S A. 116, 14260–14269. doi: 10.1073/pnas.1905985116.

Tort, A.B.L., Brankačk, J., and Draguhn, A. (2018). Respiration-Entrained Brain Rhythms Are Global but Often Overlooked. Trends Neurosci. 41, 186–197. doi: 10.1016/j.tins.2018.01.007.

Tsanov, M., Chah, E., Reilly, R., O'Mara, S.M. (2014). Respiratory cycle entrainment of septal neurons mediates the fast coupling of sniffing rate and hippocampal theta rhythm. Eur J Neurosci. 39, 957–74. doi: 10.1111/ejn.12449. Epub 2013 Dec 11.

Tyler, W.J. (2012). The mechanobiology of brain function. Nat Rev Neurosci. 13, 867–878.

Tyler,W.J., Lani, S.W., and Hwang, G.M. (2018). Ultrasonic modulation of neural circuit activity. Curr Opin Neurobiol. 50, 222–231. doi: 10.1016/j.conb.2018.04.011. Epub 2018 Apr 16.

Uhlén, M., Fagerberg, L., Hallström, B.M., Lindskog, C., Oksvold. P., Mardinoglu, A., et al., (2015). Tissue-based map of the human proteome. Science. 347, 1260419 DOI: 10.1126/science.1260419.

Urigüen, J.A., and Garcia-Zapirain, B. (2015). EEG artifact removal-state-of-the-art and guidelines. J Neural Eng. 12, 031001. doi: 10.1088/1741-2560/12/3/031001.

Usoskin, D., Furlan, A., Islam, S., Abdo, H., Lönnerberg, P., Lou, D., et al. (2015). Unbiased classification of sensory neuron types by large-scale single-cell RNA sequencing. Nat. Neuro. 18, 145–153. doi: 10.1038/nn.3881

Velasco-Estevez, M., Mampay, M., Boutin, H., Chaney, A., Warn, P., Sharp, A., et al. (2018). Infection Augments Expression of Mechanosensing Piezo1 Channels in Amyloid Plaque-Reactive Astrocytes. Front Aging Neurosci. 22, 10, 332.

Velasco-Estevez, M., Gadalla, K.K.E., Liñan-Barba, N., Cobb, S., Dev, K.K., Sheridan, G.K.. (2019). Inhibition of Piezo1 attenuates demyelination in the central nervous system. Glia. Oct 9. doi: 10.1002/glia.23722. [Epub ahead of print]

Velasco-Estevez, M., Rolle, SO., Mampay, M., Dev, K.K., and Sheridan, G.K. (2019). Piezo1 regulates calcium oscillations and cytokine release from astrocytes. Glia. 2019 Aug 21. doi: 10.1002/glia.23709.

Vinje, V., Ringstad, G., Lindstrøm, E.K., Valnes, L.M., Rognes, M.E., Eide, P.K. et al., (2019). Respiratory influence on cerebrospinal fluid flow - a computational study based on long-term intracranial pressure measurements. Sci. Rep. 9, 9732. doi: 10.1038/s41598-019-46055-5.

Wagshul, M.E., Eide, P.K., and Madsen, J.R. (2011). The pulsating brain: A review of experimental and clinical studies of intracranial pulsatility. Fluids Barriers CNS. 8:5.

Wang, J., La, J.H., and Hamill, O.P. (2019a). PIEZO1 is selectively expressed in small diameter mouse DRG neurons distinct from neurons strongly expressing TRPV1. Front. Mol. Neurosci. 12, 178.

Wang, L., Zhou, H., Zhang, M., Liu, W., Deng, T., Zhao, Q., et al. (2019b). Structure and mechanogating of the mammalian tactile channel PIEZO2. Nature. 573, 225–229.

Wang, Y.Y., Zhang, H., Ma, T., Lu, Y., Xie, H.Y., Wang W, et al. (2019c). Piezo1 mediates neuron oxygen-glucose deprivation/reoxygenation injury via Ca2-/calpain signaling. Biochem. Biophys. Res. Commun. 513(1):147–153.

Woo, S. H., Lukacs, V., de Nooij, J.C., Zaytseva, D., Criddle, C.R., Francisco, A., et al. (2015). Piezo2 is the principal mechanotransduction channel for proprioception. Nat. Neurosci. 18, 1756–1762.

Woo, S.H., Ranade, S., Weyer, A.D., Dubin, A.E., Baba, Y., Qiu, Z., et al. (2014). Piezo2 is required for Merkel-cell mechanotransduction. Nature 509, 622–626.

Yackle, K., Schwarz, L.A., Kam, K., Sorokin, J.M., Huguenard, J.R., Feldman, J.L., et al. (2017). Breathing control center neurons that promote arousal in mice. Science. 355, 1411–1415. DOI: 10.1126/science.aai7984

Yanovsky, Y., Ciatipis, M., Draguhn, A., Tort, A.B., and Brankačk, J. (2014). Slow oscillations in the mouse hippocampus entrained by nasal respiration. J. Neurosci. 34, 5949–64. doi: 10.1523/JNEUROSCI.5287-13.2014.

Yasuma, F., and Hayano, J. (2004). Respiratory sinus arrhythmia: why does the heartbeat synchronize with respiratory rhythm? Chest. 125, 683–690.

Zago, S., Ferrucci, R., Marceglia, S., and Priori, A. (2009). The Mosso method for recording brain pulsation: the forerunner of functional neuroimaging. Neuroimage. 48, 652–656.

Zeisel, A., Muñoz-Manchado, A.B., Codeluppi, S., Lönnerberg, P., La Manno, G., Juréus, A., et al. (2015). Brain structure. Cell types in the mouse cortex and hippocampus revealed by single-cell RNA-seq. Science. 347,1138–1142. doi: 10.1126/science.aaa1934.

Zelano, C., Jiang, H., Zhou, G., Arora, N., Schuele, S., Rosenow, J. et al., (2016). Nasal Respiration Entrains Human Limbic Oscillations and Modulates Cognitive Function. J. Neurosci. 36, 12448–12467; DOI: https://doi.org/10.1523/JNEUROSCI.2586-16.2016

Zeng, W.Z., Marshall, K.L., Min, S., Daou, I., Chapleau, M.W., Abboud, F.M., et al., (2018). PIEZOs mediate neuronal sensing of blood pressure and the baroreceptor reflex. Science 362, 464–467.

Zhang, M., Wang, Y., Geng, J., Zhou, S., and Xiao, B. (2019). Mechanically Activated Piezo Channels Mediate Touch and Suppress Acute Mechanical Pain Response in Mice. Cell Rep. 26, 1419–1431.e4. doi: 10.1016/j.celrep.2019.01.056.

Zhong, W., Ciatipis, M., Wolfenstetter, T., Jessberger, J. Müller, C., Ponsel, S. et al. (2017) Selective entrainment of gamma subbands by different slow network oscillations. Proc. Natl. Acad. Sci. U S A. 114, 4519–4524. doi: 10.1073/pnas.1617249114. Epub 2017 Apr 10.

